# Structural control of fibrin bioactivity by mechanical deformation

**DOI:** 10.1101/2020.08.24.265611

**Authors:** Sachin Kumar, Yujen Wang, Manuel K. Rausch, Sapun H. Parekh

## Abstract

Fibrin is a fibrous protein network that entraps blood cells and platelets to form blood clots following vascular injury. As a biomaterial, fibrin acts a biochemical scaffold as well as a viscoelastic patch that resists mechanical insults. The biomechanics and biochemistry of fibrin have been well characterized independently, showing that fibrin is a hierarchical material with numerous binding partners. However, comparatively little is known about how fibrin biomechanics and biochemistry are coupled: how does fibrin deformation influence its biochemistry at the molecular level? In this study, we show how mechanically-induced molecular structural changes in fibrin affect fibrin biochemistry and fibrin-platelet interaction. We found that tensile deformation of fibrin lead to molecular structural transitions of α-helices to β-sheets, which reduced binding of tissue plasminogen activator (tPA), an enzyme that initiates fibrinolysis, at the network and single fiber level. Moreover, binding of tPA and Thioflavin T (ThT), a commonly used β-sheet marker, was primarily mutually exclusive such that tPA bound to native (helical) fibrin whereas ThT bound to strained fibrin. Finally, we demonstrate that conformational changes in fibrin suppressed the biological activity of platelets on mechanically strained fibrin due to attenuated α_IIb_β_3_ integrin binding. Our work shows that mechanical strain regulates fibrin molecular structure and fibrin biological activity in an elegant mechano-chemical feedback loop, which likely influences fibrinolysis and wound healing kinetics.

## Introduction

Fibrin is a mesh-like, three-dimensional (3D) fibrous protein network that is the primary component of blood clots, entrapping blood cells and platelets to prevent blood loss at wound sites [1]. In its role as the fibrous network in blood clots, fibrin acts as a biochemical scaffold – binding to and interacting with various molecules that participate in the wound healing response, and as a mechanical stabilizer against forces from muscle contractions (tension) and blood flow (shear) [2]. Fibrin viscoelasticity, along with its strain stiffening behavior, have been shown to underlie haemostasis, blood clot stability, and supportive function for wound healing under the dynamic *in situ* environment [3]. These biomechanical properties of fibrin depend on its hierarchical structure, building up from the molecular to the fiber level [4]. Several studies have concentrated on understanding and evaluating structural changes in fibrin under mechanical deformation and how those changes correlate to clot stability [3-7]. In addition, Bucay *et al*. and Li *et al*. showed that stretching single fibrin fibers affected plasmin-mediated fibrinolysis, albeit with opposite effects. Bucay *et al*. showing that fiber lysis was faster for pre-strained fibers whereas Li *et al*. found that lysis was slower on strained fibrin fibers [8, 9]. Both studies hypothesized that plasmin bioactivity depends on fiber strain, however, via different mechanisms. At the network level, studies by Varju *et al*. and Adhikari *et al*. showed that tensile forces on fibrin networks reduce the degradation rate of fibrin by plasmin [10, 11]. Clearly, the role of strain in fibrin degradation is still debated, and an important element not explored in any of these studies is how fibrin degradability is sensitive to its molecular structure. Therefore, the complete picture of fibrin’s mechano-chemical response remains unknown.

In addition to degradation, variation in clot mechanical properties have been shown to affect several aspects of platelet-induced haemostasis and thrombosis [12, 13]. Platelets sense and respond to their local environment based on fibrin mechanical properties [14]. Qiu *et al*. showed that changing fibrin network stiffness modulated platelet adhesion, spreading, and activation, highlighting how a heterogeneous fibrin distribution could potentially affect platelet response [14]. Similarly, a correlation was also found between fibrin fiber diameter and platelet aggregation. Fibrin networks with thinner diameter fibers not only showed prolonged (slower) lysis, but also resulted in substantial aggregation of platelets [15]. In both of the previous examples, as in most studies described so far, the focus has been on how macroscale properties of fibrin (e.g. fiber density, fiber diameter, and fibrin network stiffness) affect platelet activity [14-17]. The molecular properties of fibrin, specifically the confirmation, or molecular structure, of fibrin monomers within the fibers, and their relation to fibrin biology and biochemistry, have remained largely absent from the fibrin structure-function literature. This is despite evidence by many groups [18-20], including us [7], showing that fibrin molecular structure changes from a dominant α-helix to a dominant β-sheet motif and that molecular distances in fibrin fibers shrink with increasing deformation [20, 21].

In this study, we relate molecular structural changes in mechanically deformed fibrin with changes in fibrin biochemistry and platelet response. Fibrin hydrogels were mechanically stretched uniaxially, and the transition of fibrin structure from α-helices to β-sheets was shown by multiple methods. Unfolding of α-helices into β-sheets upon tensile deformation resulted in enhanced binding of Thioflavin T (ThT) – a well-known β-sheet binding molecular rotor dye. The structural transition of fibrin strongly affected the binding of tissue plasminogen factor (tPA), an enzyme that converts plasminogen to plasmin upon fibrin binding, showing evidence of altered fibrin biochemical activity upon tensile deformation. In addition, platelet activity was strongly suppressed on mechanically deformed fibrin due to structural alterations that decreased α_IIb_β_3_ integrin binding to RGD motifs. This work provides a mechanistic perspective on how mechanical strain on fibrin affects not only fiber morphology, fiber density, and orientation, but also constituent protein structure at the molecular level, regulating the biological activity of fibrin.

## Materials and Methods

### Preparation of fibrin hydrogel substrate

**Figure 1** shows the preparation process for the fibrin hydrogels preparation in our experiments. In brief, a thick (1 mm) PDMS sheet (Gel-Pak) was cut into a rectangular mold (PDMS mold) and placed on a supporting, thin PDMS sheet (100 µm thick) (Gel-Pak) that was on a detergent (Micro-90) cleaned glass slide. The whole assembly was treated with ultraviolet ozone (UVO) for 30 min to render the surfaces hydrophilic. Thereafter, a fresh fibrinogen solution (20 mM HEPES, 5 mM CaCl_2_ and 150 mM NaCl at pH 7.4; fibrinogen (FIB 2) 7.5 mg/ml and thrombin (Factor IIa) 1.0 U/ml (Enzyme Research Laboratories)) [7] was prepared, filled into the hollow rectangular cavity, and covered with another thin PDMS sheet (non-UVO treated) in direct contact with the solution and a coverslip. The sandwiched fibrinogen solution was allowed to polymerize by incubating at 37°C for 2 hr. After polymerization, the covering thin PDMS sheet and coverslip and rectangular PDMS mold were carefully and slowly removed to obtain an intact fibrin hydrogel on the supporting PDMS sheet.

**Figure 1:**
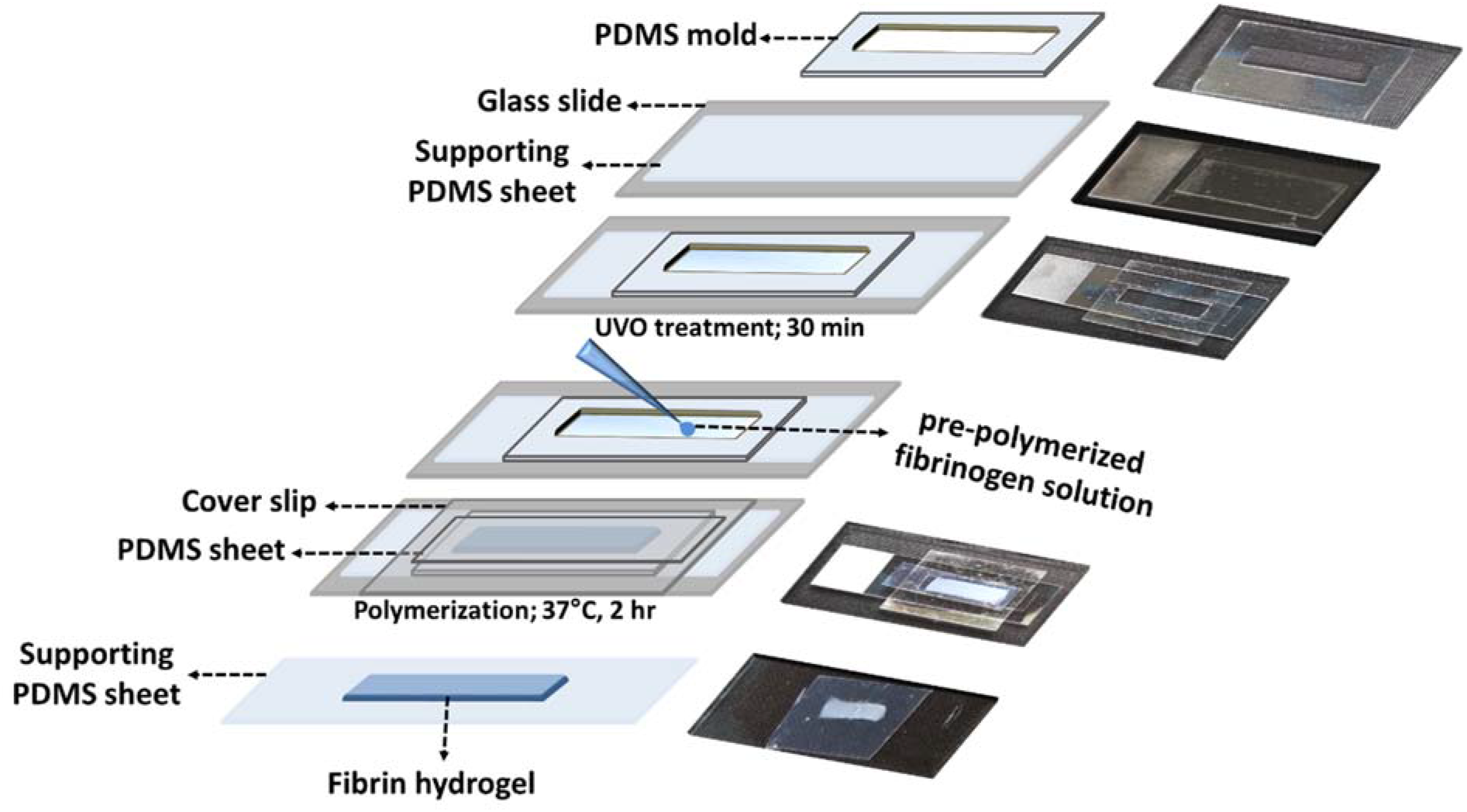
Schematic illustration fibrin hydrogel and PDMS substrate preparation. Digital images show the actual process of sample assembly.

### Tensile deformation of fibrin gels

For tensile deformation of fibrin networks, the thin supporting (100 µm thick) PDMS sheet upon which the polymerized fibrin gel adhered, was stretched with a motorized tensile stretcher to 100% of its initial length, thereby also stretching the fibrin gel, as shown in **Movie S1 and Supporting Figure S1(i)**. Thereafter, the stretched fibrin gels were clamped using in-house clamps as shown in **Figure S1(ii)** to hold the strain on the supporting PDMS sheet for further studies and characterization. All studies and characterization on clamped fibrin samples were performed within hours, and the sample was hydrated fully the entire time.

### Characterization of fibrin morphology

Fibrin morphology was imaged using an upright laser scanning confocal microscope (LSM-510, Zeiss) under reflectance using a 40X, 1.0 NA water-dipping objective (Zeiss). Fibrin network samples were illuminated with a 488 nm laser and reflected light was collected by setting the Meta detector channel between 475-500 nm wavelength in accordance to previous literature [22]. The diameter of fibrin fibers was measured on raw images using ImageJ. In addition, AC mode atomic force microscopy (AFM, Dimension Icon, Bruker) was used to evaluate morphology of fibrin sample under unstrained and strained conditions.

### Molecular structure characterization of fibrin

Changes in fibrin molecular structure upon mechanical deformation were evaluated using attenuated total internal reflection infrared (ATR-IR) spectroscopy measurements. For these experiments, unstrained and tensile strained fibrin networks were mounted on custom-built hollow Teflon molds (shown in **Figure S1(iii)**) and placed over the ATR crystal surface to achieve good contact. ATR-IR spectra were collected using a TENSOR II Bruker spectrometer equipped with a DGTS detector. Typically, for each spectrum, 128 interferograms were collected and averaged in single-beam mode at a resolution of 4 cm^-1^. Spectra from three different spots on each sample were averaged for analysis. For structural analysis, decomposition of the ATR-IR spectra of the Amide I and II regions using OriginPro was done by setting component peak locations found by second-derivative analysis (**Figure S2**) in accordance to previously reported literature [19]. Lorentzian profiles were used to fit the envelope of the amide band spectra. The peak widths and their positions were assigned to respective secondary structure in accordance with previous work on fibrin [19, 23] (detailed information is provided in the Supporting Information). Peak areas were estimated by using least-squares curve-fitting procedures in OriginPro to quantify the fraction of respective secondary structures in protein.

### Imaging biochemical activity and molecular structure of molecular fibrin

Fluorescent species (Thioflavin T (ThT, Sigma)), and tissue plasminogen activator (tPA) with a FITC label (tPA-FTIC, Abcam) were used to probe the molecular structure and biochemistry of fibrin, respectively. A ThT solution (100 µM) was prepared as specified by the supplier and incubated with fibrin networks (both unstrained and strained) for 30 min. After multiple washes with phosphate-buffered saline (PBS), bound ThT fluorescence was imaged by exciting at 450 nm and collecting emission from 480-530 nm. Similarly, fibrin networks (unstrained and strained) were incubated with a tPA-FITC (20 µM) protein solution for 30 min, washed with PBS multiple times, and imaged in fluorescence by exciting at 488 nm and collecting emission from 530-580 nm. All imaging for these experiments used the LSM-510 confocal microscope with the same objective as described above, and the settings as described previously [22].

### Platelet isolation and measurement of attachment and morphology on fibrin hydrogels

For platelet isolation, blood from a healthy volunteer was drawn into acid citrate dextrose (ACD) and EDTA vacutainer tubes by a licensed physician in accordance with the standard protocol approved by the institute. Platelets were isolated from whole blood by differential centrifugation in accordance with the previously established methods [24]. In brief, the whole blood sample was centrifuged at 800 rpm for 10 min to obtain platelet rich plasma (PRP). PRP was then centrifuged at 1500 rpm for 5 min to obtain a platelet pellet. The platelet pellet was suspended and diluted to final platelet density of 10^5^ per ml in HEPES Tyrodes buffer (pH 7.4) supplemented with 0.1% BSA. The morphology of the obtained platelets was verified under brightfield microscopy (IX-81, Olympus) with a 40X, 0.75 NA objective (Olympus).

The obtained platelets were immediately labeled with 1µM Calcein green AM dye for 15 min at room temperature and seeded on unstrained or strained fibrin hydrogels as done previously [25]. The platelet-fibrin mixture was incubated for 2 hr at 37°C in CO_2_ incubator. After incubation, samples were washed several times with HEPES Tyrodes buffer to remove non-adhered platelets. The remaining adhered platelets were fixed with 4% formaldehyde solution. Platelet attachment and morphology were quantified using fluorescence with the LSM-510 confocal laser scanning microscope and scanning electron microscopy (SEM, LEO Gemini 1530, Germany). All platelet experiments were performed within 5 hr of whole blood collection to avoid cell death and intrinsic activation.

### Binding assay of platelet integrins to fibrin

Platelet integrin (α_IIb_β_3_) binding activity to fibrin was tested using microspheres coated with the extracellular domain of α_IIb_β_3_ integrin. Recombinant human α_IIb_β_3_ integrin (Abcam) (1 mg/ml) and fluorescent microbeads (Fluoresbrite-Carboxylate, 1.75μm diameter) suspended in PBS (10^8^ beads/ml) were mixed, and the solution was then placed on a shaker at 300 rpm for 24 hr at 4°C for adsorption of integrins onto beads. Beads were collected by centrifugation (10000 rpm for 5 min) and washed twice with PBS to remove unbound integrin. Thereafter, integrin-coated beads were blocked with 2 % BSA-FITC (Sigma) solution for 24 hr at 4°C. For control experiments, BSA-coated beads were prepared in accordance to above mentioned protocol without any integrin coating. Finally, integrin-coated (or BSA-coated) beads were pelleted, washed twice, and re-suspended in PBS (10^7^ beads/ml) having 1 mM MnCl_2_ (or just PBS for BSA-coated beads). SDS-PAGE was used for characterization of bound proteins (integrin α_IIb_β and BSA) to the microbeads **(Figure S4)**.

For the integrin binding assay, unstrained and strained fibrin hydrogels were incubated with integrin-coated (or BSA-coated) microbeads and allowed to bind for 1 hr at room temperature followed by multiple rinses with PBS to remove unbound beads. Rinsed fibrin samples were imaged using confocal laser scanning microscopy (LSM 510), and the bead density was measured manually from multiple fields of view using ImageJ. After averaging bead density for α_IIb_β_3_-coated beads from multiple fields of view, the density of BSA-coated beads bound to similar fibrin hydrogels was subtracted to account for any non-specific interaction.

### Statistical analysis

Significant differences between experimental groups were analyzed using 1-way ANOVA (analysis of variance) with Tukey’s test for multiple comparisons using OriginPro. Differences were considered statistically significant for p < 0.05 and indicated by symbols in the figures.

## Results

Thin PDMS sheets, upon which fibrin networks were polymerized, were initially treated with ultraviolet ozone (UVO) to render them hydrophilic. UVO treatment to PDMS has been well established to increase surface wettability and adhesion properties of PDMS for protein adsorption [26] [27]. As our goal was to investigate the structure-function relation of fibrin under tensile loads, UVO treatment ensured that the fibrin network strongly adhered to the PDMS sheet to which we applied tensile strain. **Figure 1** shows that the prepared PDMS-fibrin construct not only supported the fibrin network, but also facilitated mechanical deformation of fibrin under tension (shown in **Movie S1 and S2**). We applied 100% strain to the flexible PDMS sheet, and this was transmitted to the fibrin network; hereafter called 100% strained or strained fibrin. Tensile deformation of fibrin clearly showed distinct morphological changes in terms of fibrin fiber orientation, fiber diameter, and fiber packing density as shown by confocal reflectance images (**Figure 2(i)**).

**Figure 2:**
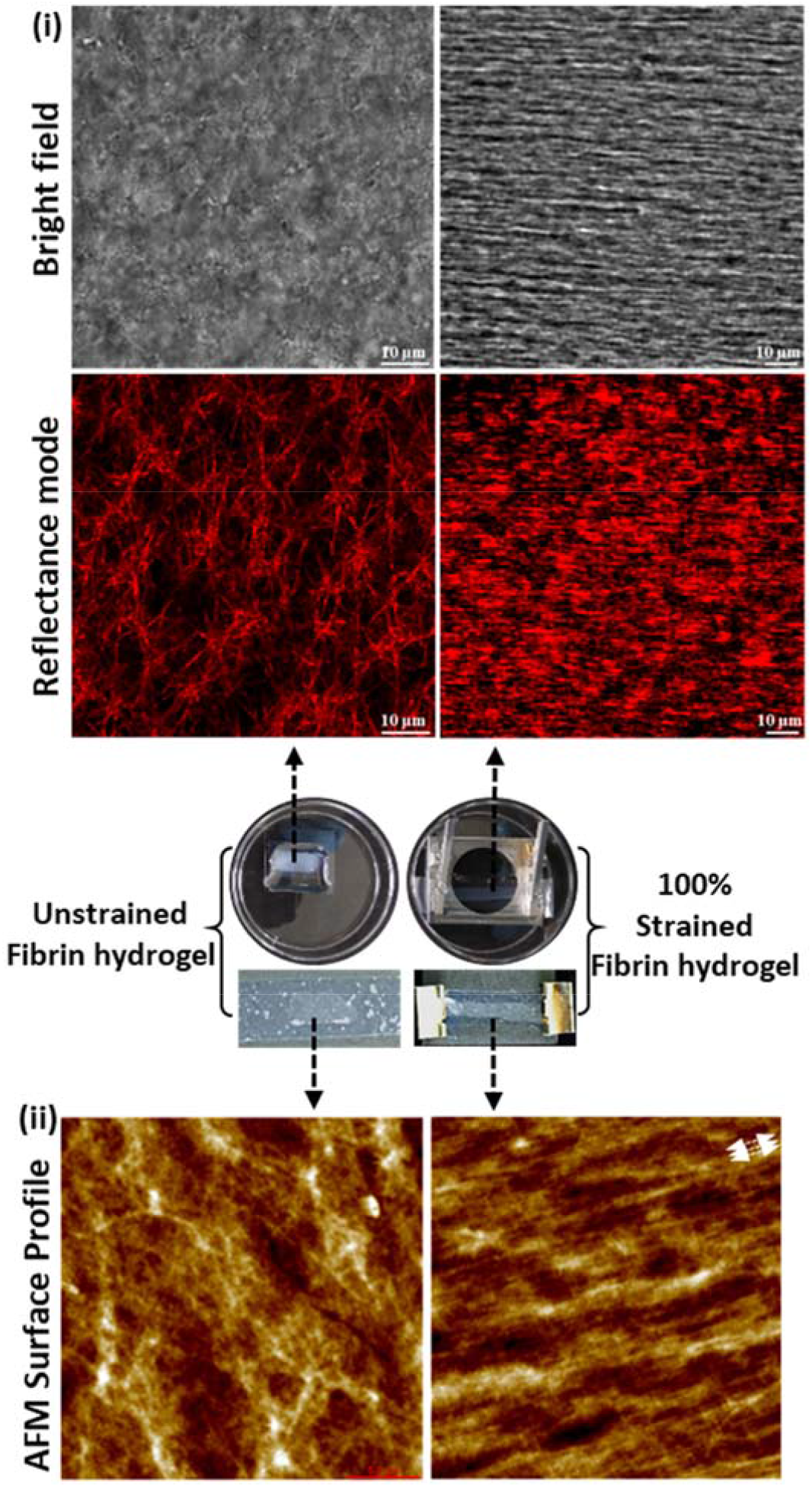
Morphology change of fibrin under mechanical deformation. (i) Transmitted light (top) and reflectance (bottom) confocal micrographs of unstrained and 100% strained fibrin networks. Strain was applied in the horizontal lab frame. Images are taken ∼ 50 µm deep into network from the free (top) surface. (ii) AFM surface profile of unstrained and 100% strained fibrin (white arrows indicate the stretching direction). The middle photos show samples corresponding to unstrained and 100% strained fibrin samples.

Unstrained (as prepared) fibrin hydrogels showed an isotropic fibrous network structure with an average fiber diameter of 0.4 ± 0.1 µm (mean ± standard deviation, N > 50) as measured from the confocal reflectance images. Upon straining, fibrin fibers elongated, with the network expelling water and thinning; the fiber diameter in strained gels decreased to 0.25 ± 0.05 µm (N > 50), as similarly reported by others [10, 18]. In addition, fibers in 100% strained fibrin aligned strongly in the stretching direction and came into close contact with one another, forming a densely packed fiber structure that nearly masked the borders of individual fibers. Studies have previously reported that elongation and alignment of fibers under mechanical strain indicates a transition from a soft, randomly oriented network architecture to a packed and aligned structure that results in a stiffer network due to strain-stiffening of fibrin fibers [10, 28]. Similar to confocal reflectance, AFM images of the free (top) fibrin surface of unstrained fibrin hydrogels displayed randomly oriented fibers while the stretched fibrin hydrogel showed elongated and aligned fibers in the stretching direction (indicated by white arrows) with densely packed structures (**Figure 2(ii)**).

Mechanical deformation of fibrin not only influences fiber scale morphology but also affects molecular structure [3, 7]. Fibrin molecular structural changes due to mechanical deformation have been studied using vibrational spectroscopy, which provides valuable information on protein secondary structure (α-helix, β-sheet, turn, and random coil structures)[29].

**Figure 3(i)** shows ATR-IR spectra from 1200 to 1750 cm^-1^ for unstrained and 100% strained fibrin. The spectra show characteristic amide peaks (∼ 1650 cm^-1^ for Amide I, ∼ 1550 cm^-1^ for Amide II band, and ∼ 1250 cm^-1^ for Amide III). Differences in the amide band shape were observed upon stretching, especially for the Amide I peak. The fibrin Amide I peak position for 100% strained fibrin shifted to lower frequency (1628 cm^-1^) compared to unstrained fibrin (1645 cm^-1^) (**Figure 3(i)**), indicative of increased β-sheet content after stretching. We used second derivative analysis (**Figure S2)** and subsequent spectral decomposition of the Amide I and II regions to provide quantitative information on the secondary structure before and after strain application (**Figure 3(ii), Table S1 and S2**). This analysis showed that applying tensile forces to fibrin resulted in a 36 % increase in β-sheet content, whereas α-helix content decreased by 31% compared to the unstretched fibrin structure. Our results are quantitatively consistent with our previous study and in good agreement with literature, showing that tensile deformation of fibrin tension is accompanied by α-helix to β-sheet molecular structure transitions [7, 19, 30, 31].

**Figure 3:**
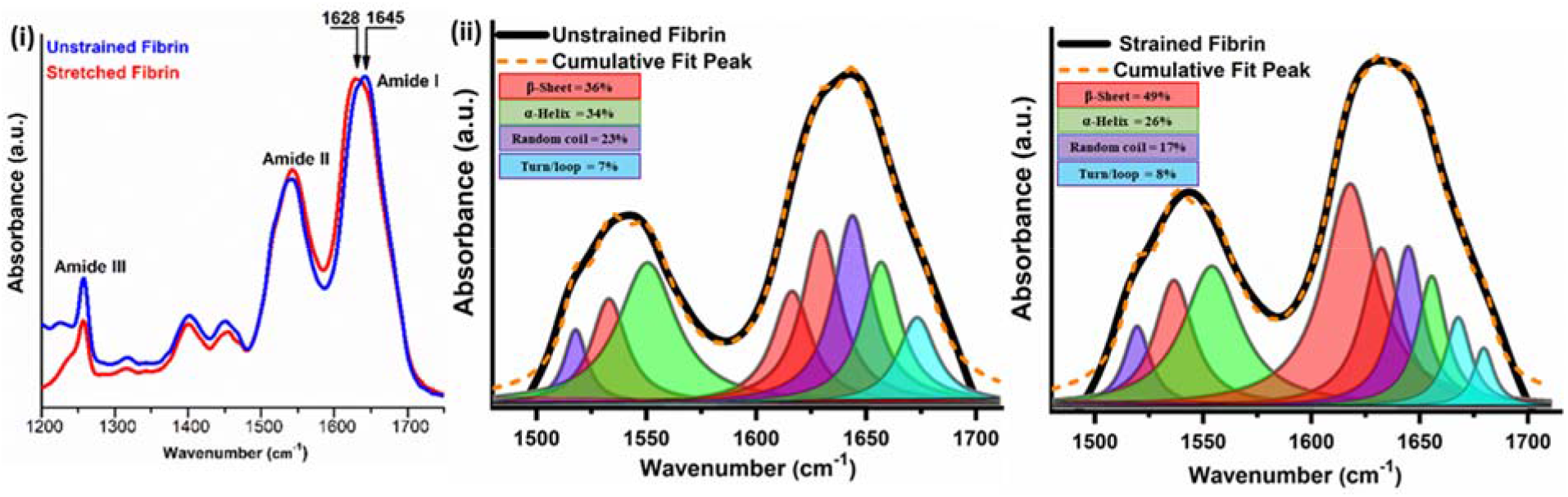
Tensile strain causes fibrin molecular structural unfolding. (i) ATR-IR spectra of unstrained and 100% strained fibrin showing Amide I to III bands with a clear Amide I peak shift showing structural changes. (ii) Decomposition of Amide I and II modes into constituent peaks corresponding to different protein secondary structures shows the increased β-sheet content with strain.

Our goal was to investigate if the molecular structural changes in fibrin modulate its biochemical activity. Thus, we moved forward to determine if relaxed or strained fibrin exhibited differential biochemical activity for two molecules: Thioflavin T (ThT) and tissue plasminogen activator (tPA). ThT is a so-called molecular rotor dye, which has been extensively used as a standard biophysical probe for detecting β-sheet rich protein structures [32, 33]. Binding of ThT to β-sheet domains has been shown to increase the fluorescence yield (and decrease the fluorescence lifetime) of ThT [34].

Strained fibrin networks showed a significant increase in ThT fluorescence intensity compared to native, unstretched fibrin hydrogels (**Figure 4a (i)**), suggesting a more β-sheet rich environment in fibrin after loading, in agreement with our spectroscopy results above. In contrast, ThT fluorescence was substantially less pronounced in relaxed fibrin gels under the same imaging conditions. Relaxed fibrin exhibits primarily α-helices in the coiled coil region and some parallel β-sheet content in the D domains, even in the native state, based on the fibrinogen crystal structure. With the substantially weaker ThT signal in unstrained fibrin gels, we surmise that strong ThT fluorescence in 100% strained fibrin came from increased ThT interaction in the coiled-coil region that becomes β-sheet rich under tensile strain [30].

**Figure 4:**
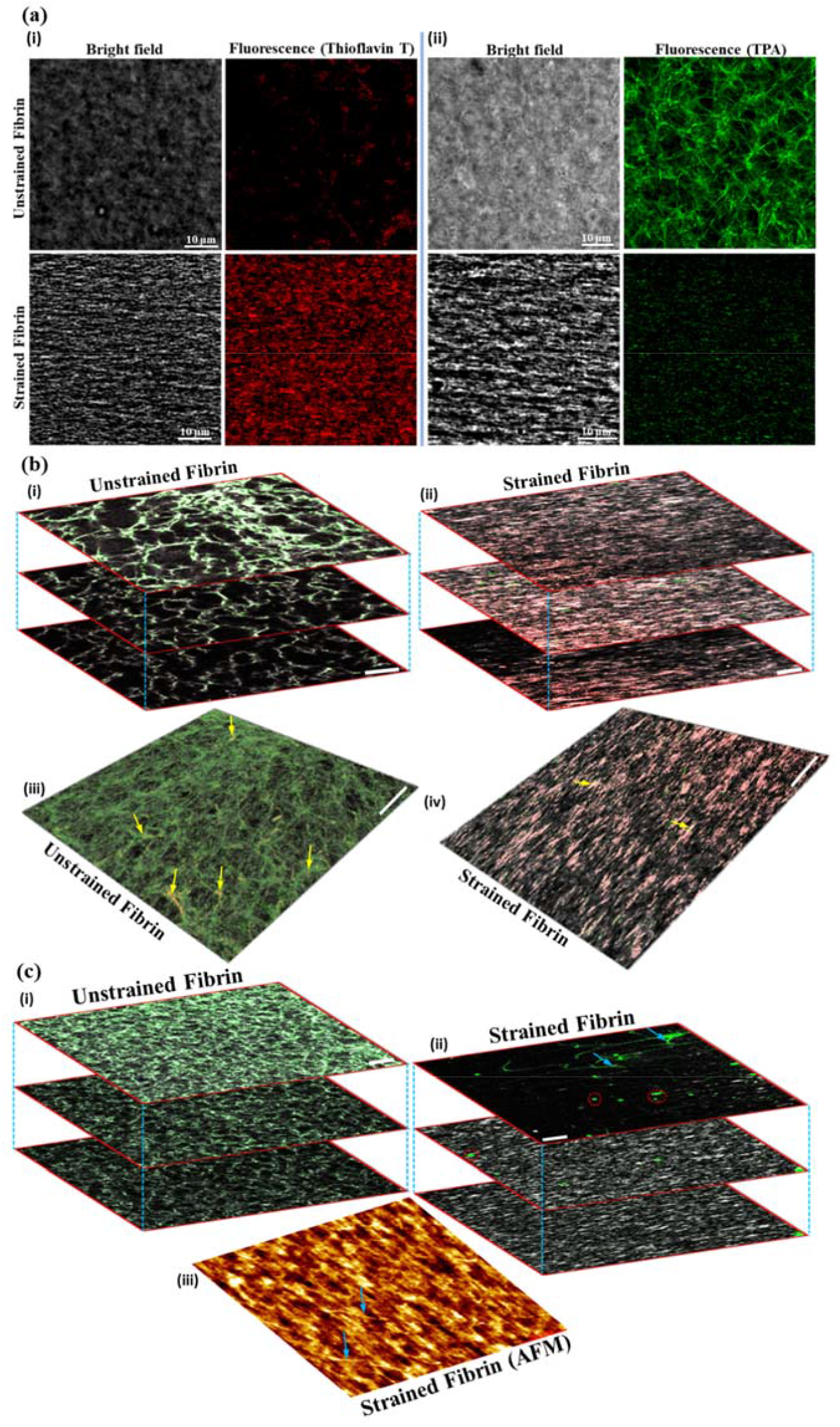
Mechanical deformation of fibrin affects ThT and tPA binding to fibrin. (a) (i) Binding of ThT on unstrained and 100% strained fibrin. (ii) Binding of tPA on unstrained and 100% strained fibrin. The imaging settings were identical for ThT images in (i) and for tPA images in (ii), as are the display contrasts. (b) Differential binding and distribution of tPA and ThT to unstrained and 100% strained fibrin. (i) and (ii) Images at different sample depths (top to bottom: surface to 50 µm depth) profile of tPA (green) and ThT (red) bound to unstrained and 100% strained fibrin (grey), respectively. (iii) and (iv) Z-projected images (surface to 50 µm depth) of co-localization of tPA (green) and ThT (red) on unstrained and 100% strained fibrin networks, respectively. Yellow arrows in shows areas of ThT and tPA co-localization. (c) (i) Images at different sample depths (top to bottom: surface to 50 µm depth) showing tPA binding (green) on unstrained fibrin (grey). (ii) Images at different sample depths (top to bottom: surface to 50 µm depth) of the same fibrin sample as in c(i) shows unbinding of tPA upon 100% tensile deformation. Red circles show highlight broken fibers with tPA binding after straining fibrin. (iii) AFM profile of tensile deformed fibrin sample. Arrows show broken fibers. Scale bar = 10 µm for optical images in b and c.

During the wound healing process, fibrin must also be enzymatically degraded as the blood clot is slowly replaced with new tissue. In this capacity, fibrin also serves as a biochemical substrate for plasminogen cleavage by tPA that is bound to fibrin. Because strained fibrin networks showed changes in secondary structure, we hypothesized that tPA binding would be sensitive to the tensile state – and by extension the structure – of fibrin. tPA binding to fibrin has been reported to depend on fibrin structure via specific amino acids in coiled-coil region (residues 148-160 in the Aα chain) and in the D region of γ chain (residues 312-324) [35-38]. Upon incubation with tPA, unstrained fibrin showed effective binding to fibrin fibers, with tPA essentially illuminating the fiber network structure of the network (**Figure 4a (ii)**). On other hand, strained fibrin exhibited significantly lower fluorescence when incubated with tPA.

In order to demonstrate the structural specificity of tPA and ThT binding, ThT and tPA solutions were added simultaneously and allowed to bind on unstrained or strained fibrin hydrogels. The z-stack projections (see **Movie S3 and S4**) in **Figure 4b (i) and 4b (ii)** show depth distribution profiles of bound tPA (green) and ThT (red) in unstrained and 100% strained fibrin samples, respectively. ThT and tPA not only bound to the free surface but also were able to diffuse into the gel and bind to fibrin fibers at depth of 50 µm. ThT and tPA both bound to fibrin, but did so under different conditions, as suggested by the data in **Figure 4a**. ThT binding was much more prominent on strained fibrin while tPA was more prominent on unstrained fibrin. We note, however, that the distribution of both tPA and ThT in depth was more uniform and produced stronger fluorescence deeper within unstrained fibrin compared to strained fibrin networks, consistent with previous work [11]. In addition, the Z-projected images of unstrained fibrin (**Figure 4b (iii)**) showed very few spots of colocalization (indicated by arrows) between tPA (green) and ThT (colored in red). The colocalization between tPA and ThT was virtually nonexistent for 100% stretched fibrin (**Figure 4b (iv)**). These experiments indicate that tPA and ThT bind primarily in mutual exclusion and in a structure-dependent fashion, where tPA binds to fibrin adopting a more helical conformation and ThT binds to fibrin adopting a more β-sheet conformation.

In order to resolve the question of whether limited mobility within the fibrin network or the structural change of fibrin was responsible for decreased tPA binding, we performed experiments to determine if bound tPA was able to unbind and diffuse out of the gel after strain application. tPA was initially bound to an unstrained fibrin hydrogel (**Figure 4c (i)**), and upon stretching the gel, showed significantly decreased fluorescence after washing (**Figure 4c (ii)**, see **Movie S5** for more details), even at a depth of 50 µm into the network. We note that the surface of stretched fibrin, showing mostly broken or partially strained fibers retained some tPA binding (indicated by red circles in (**Figure 4c (ii)**) whereas intact and stretched fibers in the gel interior showed nearly complete loss tPA fluorescence. Therefore, binding of tPA to unstrained fibrin was reversed by tensile deformation, and the unbound tPA molecules could be subsequently washed away. This experiment shows that tPA mobility out of (and by analogy into) fibrin gels (with a [fibrinogen] = 7.5 mg/mL) is not the rate-limiting step for tPA binding to fibrin. As further evidence for reduced tPA binding to strained fibrin, we evaluated binding and unbinding of tPA on single fibrin fibers with mechanical deformation. In these experiments, only a few individual fibers were present, and all fibers were completely submerged in an aqueous environment. Consistent with the results at the network level (**Figure 4**), 100% tensile deformation of individual fibers clearly showed a significant decrease in fluorescence intensity for bound tPA molecules (**Figure S3)**. The decreased tPA signal on individual, stretched fibers further confirmed that strained fibrin has a lower affinity for tPA, which is independent of fibrin network molecular mobility. Rather, binding of tPA to fibrin likely depends on the structure of fibrin, which can be modulated with mechanical deformation that causes fibrin unfolding at the molecular level from α-helices into β-sheets. This structure-dependent force accommodation mechanism thus strongly reduces binding of tPA molecules to fibrin and therefore likely affects fibrin degradability.

In order to further evaluate the biological activity of mechanically deformed fibrin, we studied platelet response on unstrained and strained fibrin gels. Platelets were isolated from a healthy individual blood donor as described in the methods section and were well-dispersed without any aggregates, indicating no activation, at the time of our experiments (**Figure S5**). Platelets on fibrin samples showed different features, depending on if they were incubated on strained or unstrained gels. Platelets showed nearly 4-fold more attachment and more aggregation on unstrained fibrin compared to strained fibrin hydrogels (**Figure 5(i)-(iii) and Figure S6 (i)-(iii)**). In addition to differential attachment and aggregation, platelets on unstrained fibrin showed more spreading with nearly 35% of platelets having multiple cytoplasmic projections (on average ∼3 projections/per platelet). Platelets on 100% strained fibrin by comparison were round with fewer cytoplasmic projections (**Figure S6 (iv) and (v)**).

**Figure 5:**
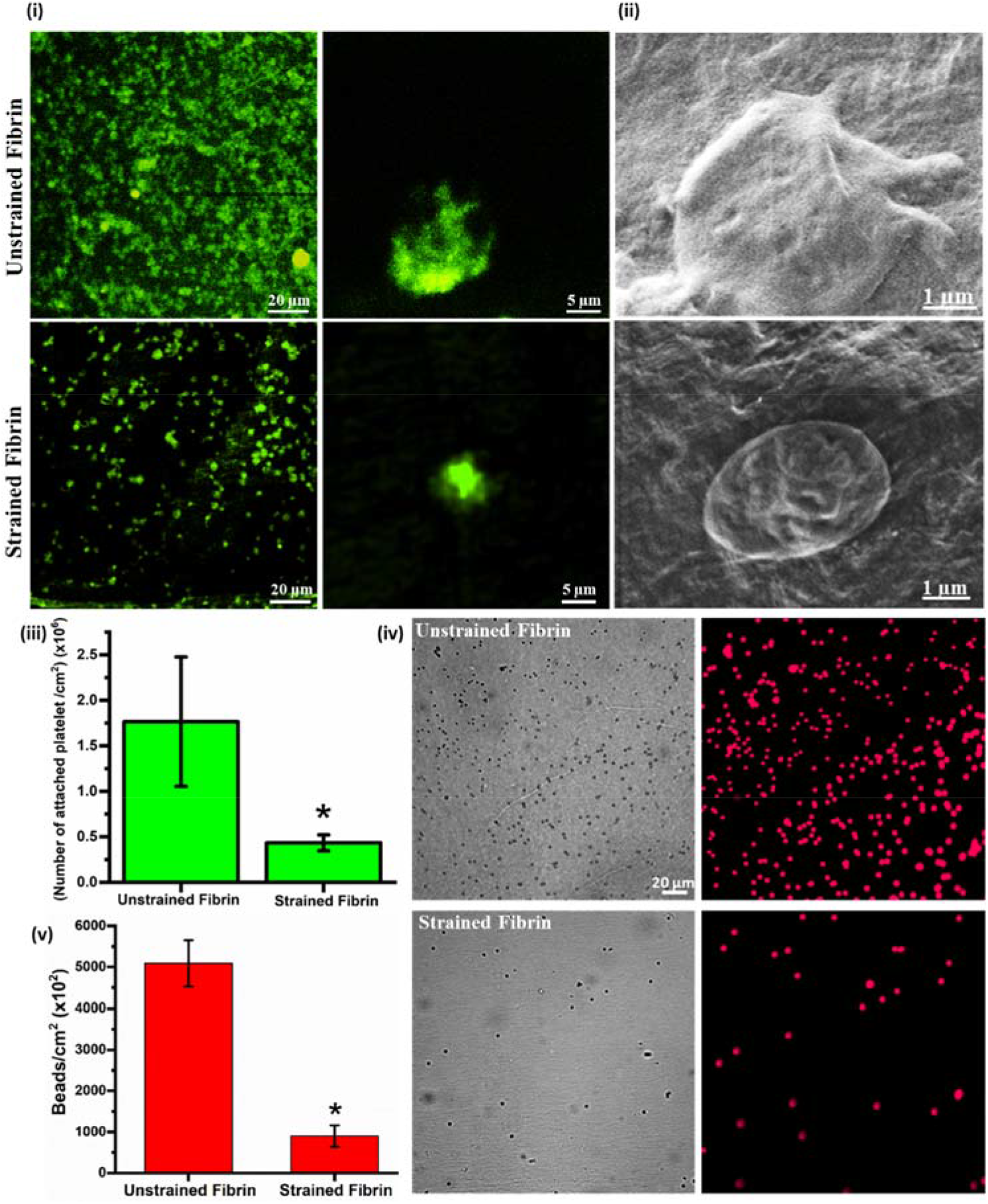
Attachment, morphology and α_IIb_β_3_ integrin binding of platelets on different fibrin surfaces. (i) Fluorescence micrographs of attached platelets stained with Calcein on unstrained and strained fibrin gels. (ii) SEM micrographs of platelets on unstrained and strained fibrin showing distinct morphology. (iii) Quantification of attached platelets on unstrained and strained fibrin. (iv) Fluorescence micrograph of bound integrin-coated micro-beads on unstrained and strained fibrin. (v) Quantitative plot of bound integrin coated micro-beads on unstrained and strained fibrin. Statistically significant differences (p<0.05) compared to unstrained fibrin is indicated by ‘*’.

Cytoplasmic spreading, aggregation, and formation of multiple projections on unstrained fibrin indicated that platelets were in an activated state [39]. We surmise that platelets adhered to strained fibrin were not activated as they showed limited spreading and no obvious projections.

Platelet adhesion and activation are regulated through the interplay of adhesive receptors and ligands. Ligand binding to cell surface integrins, specifically α_IIb_β_3_, mediates platelet adhesion, spreading, granule secretion, aggregation, and clot retraction [40]. Platelet integrin α_IIb_β_3_ has been shown to interact with γAGDV (residues position at 409 in the γ chain), αRGDF and αRGDS (residues positions 95 and 572, respectively, in the αA chain) sites on fibrinogen and fibrin [41-43]. Interestingly, Litvinov *et al*. found that integrin α_IIb_β_3_ binding was strong and significantly increased on fibrin Aα chain RGD motifs compared to variants of fibrin lacking Aα chain RGD [44]. Given that we saw changes in platelet adhesion and activation with fibrin strain, where the structure of the Aα coiled-coil regions showed structural transitions, we measured if platelet integrin binding was modulated by stretching fibrin. Microbeads coated with the extracellular domain of integrin α_IIb_β_3_ and were incubated on unstrained and 100% strained fibrin hydrogels to test for integrin binding. After several washes, we imaged the attached beads and found that integrin α_IIb_β_3_ coated beads attached more than 5-fold more on unstrained fibrin in comparison to strained fibrin (**Figure 5(iv) and (v)**). This result shows that molecular structural changes in fibrin with tensile deformation directly affected integrin engagement, therefore influencing platelet adhesion and activation.

## Discussion

Our results show that pulling on fibrin with near 100% tensile strain results in structural unfolding of the constituent proteins that make up the network, which ultimately regulates various biochemical interactions, including platelet interaction. tPA binding to fibrin is critical for clot removal, and tPA binds primarily to the coiled-coil region (and to a lesser extent to the nodule domains). Our data suggest that unfolding the coiled-coil from helices into sheets, by tensile strain, occludes the binding site in the coiled-coil region. Together with previous work showing how fibrin degradation is reduced with tensile strain, we suggest a structure-function connection between the binding of tPA (and fibrin degradation) and fibrin blood clot stability. The structure-based binding of tPA and the structural transition of fibrin under load forms an elegant mechano-chemical feedback loop that is intrinsically regulated, as we describe below. Only fibrin fibers that contain natively folded fibrin monomers, and thus not bearing large loads, are subject to degradation. Other loaded fibers in the gel are “marked” as load bearing by structural changes from helices to sheets, and the binding of tPA, and thus degradation, is drastically reduced. The ultimate outcome of such a mechanism is that the parts of the fibrin clot that are actively working (load bearing) are retained while those not load-bearing are available for degradation and further processing via the wound healing cascade. This idea with consistent with the “use it or lose it” module proposed by Dingal and Discher for intracellular cytoskeletal networks [45, 46].

Studies have shown that platelets attach to fibrin during blood clotting using the α_IIb_β_3_ integrin in a mechanosensitive way. Platelets have been shown to sense anisotropic fiber orientation and network heterogeneity by applying piconewton (pN) forces [24, 47] via cytoplasmic, pseudopodial projections. The fact that platelet binding and activation are also regulated by tensile load to fibrin clots, suggests a similar mechano-chemical feedback loop between platelets and fibrin as with tPA and fibrin. Activated platelets entrapped within a fibrin mesh exert significant force (ranging from 1.5 - 70 nN) on fibrin [24]. According to recent work showing that platelets interact with and pull on 5-10 fibers in a fibrin clot [25], the force per fiber is easily above the level required to unfold the helical coiled-coil region of ∼ 75 pN [18, 24, 48, 49]. Thereby heterogenous platelet contraction of soft fibrin clots into highly packed and stiffer clots, can modify the molecular structure of fibrin during contraction (by pulling against other contracting platelets or attachment points in the clot). As the RGD binding site of platelet integrins is located in the coiled-coil region, structural changes, e.g. secondary structure unfolding, could reduce the binding efficiency and activation of additional platelets to the contracted clot. This idea consistent with experimental work from Lishko *et al*., who showed that nascent fibrin gels supported platelet attachment, but the same gel, after removing attached platelets, showed substantially reduced platelet attachment. Their result suggests that the initial platelet adhesion altered fibrin in a way that reduced subsequent adhesion [50]. Our results show that tensile mechanical deformation of fibrin alters the molecular structure of fibrin, including in the coiled-coil region, which drastically lowered platelet activation and binding, as well as binding of the α_IIb_β_3_ integrin. Therefore, we suggest that platelet contraction of fibrin with sufficient force could alter the structure of the integrin binding domains (RGD motifs) on Aα chains, affecting secondary interaction or adhesion of platelets on strained fibrin networks.

Our proposed mechano-chemical platelet fibrin interaction is summarized in **Figure 6**. Sufficiently large tensile mechanical deformation of fibrin alters the structure of fibrin with densely packed, stiff, and aligned fibers. When the deformation is large enough, the molecular structure of fibrin proteins changes from α-helices to β-sheets in the coiled-coil region as indicated in step 1. As the RGD binding site of platelets is located in the Aα coiled-coil region, secondary structural changes can reduce its accessibility or binding compatibility. Thus, the interaction between α_IIb_β_3_ the RGD binding site on fibrin is attenuated (step 2), and because the α_IIb_β_3_-RGD interaction is mechanosensitive, platelets sense insufficient resistance from the fibrin network that is required to form strong interactions and trigger activation. On the other hand, platelets engaging with unstrained fibrin bind strongly to the RGD domain on natively folded coiled-coils. This provides strong interaction and adhesion forces for attachment, which results in more platelet spreading and activation showing multiple cytoplasmic projections (step 3). Thereby, molecular structural changes in fibrin can affect biochemical and biological activity and highlight the unique mechano-chemical properties of fibrin.

**Figure 6:**
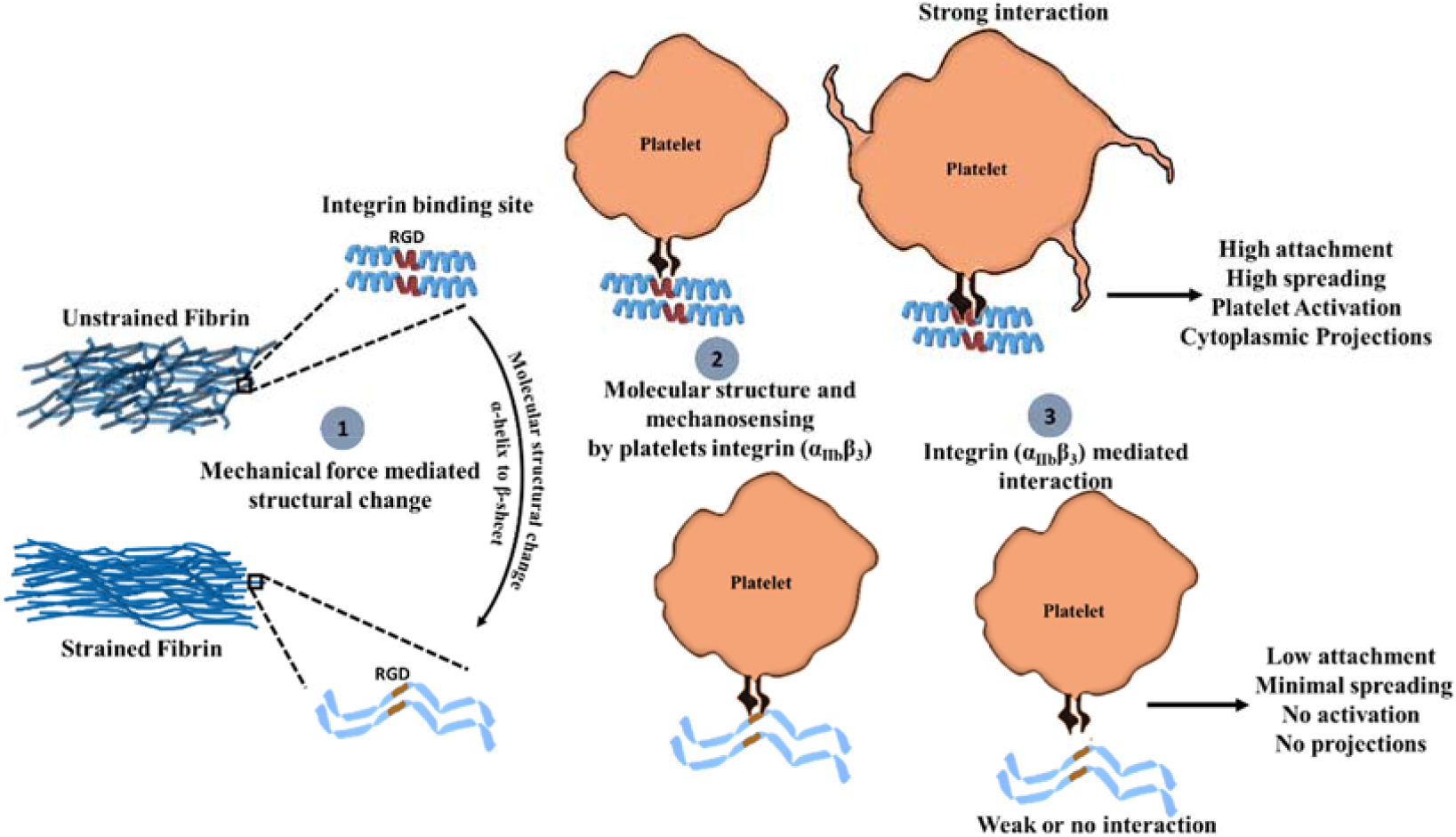
Proposed mechanism for mechano-chemical platelet activation on fibrin.

While our work shows strong evidence for structural transitions at the molecular level in fibrin being responsible for modifying biological activity of fibrin, we cannot completely exclude the possibility that squeezing water out of the intra-fiber or inter-fiber space could alter tPA binding or platelet attachment. Weisel and colleagues have shown that the average distance between the protofbrils is ∼ 5 nm under well-hydrated conditions, suggesting that macromolecules like tPA and plasminogen having diameters ∼ 6 nm (based on molecular weight) are already somewhat excluded from the intra-fiber space even before stretching [51]. In addition, the data in **Figure 5(iv) and (v)** on integrin binding suggests that increased packing contributes to a lesser extent than structural transitions. The distribution of extracellular integrin domains on the microbeads can be assumed to be uniform, and the density of RGD binding sites on fibrin would only increase with tensile loading (due to fiber packing). Therefore, decreased microbead binding to stretched fibrin, which has undergone structural changes where the primary RGD binding site exists, suggests that platelet attachment (and activation) is decreased because of structural changes rather than fiber packing.

## Conclusions

The experiments presented here show that molecular structural changes in fibrin affect biochemical and biological activity, highlighting a unique mechano-chemical sensitivity of fibrin biomaterials. Fibrin strained to near 100% in tension showed densely packed and aligned fibers with secondary structural changes (α-helices to β-sheets) that prevented integrin-mediated platelet adhesion and activation as well as enzyme (tPA) binding. While our results demonstrate the concept of mechano-chemical regulation of by fibrin using two extreme cases, 0% and 100% tensile strain, clearly a subtler behavior exists. The features provided by such a mechano-chemical mechanism exemplify a structure-based response that is tuned for particular biophysical functions. The sensitivity of platelet attachment and activation on fibrin suggests that platelets sense fibrin fiber anisotropy, mechanical stiffness, and molecular structure. While we concentrate on fibrin, we note that other biomaterials, such as collagen, also show similar effects [45], suggesting that mechano-chemical regulation of biomaterials is a broad phenomenon and could possibly be exploited for biomedical applications.

## Acknowledgements

We thank Florian Gericke and Marc-Jan van Zadel for technical support and Sabine Pütz for laboratory support. S.K. acknowledges Alexander von Humboldt Foundation Postdoctoral Fellowship, and S.H.P acknowledges support from the Welch Foundation (F-2008-20190330), Texas 4000 funding, and the Human Frontiers in Science Program (RGP0045/2018).

