## Supporting Information for "Structural control of fibrin bioactivity by mechanical deformation"

**This supporting information file includes**

**Supporting Figures and data**

**Figure S1**: Fibrin stretching on thin PDMS sheets

**Figure S2**: Second derivative ATR-IR spectra of unstrained and 100% strained fibrin

**Table S1:** Fitting parameters for ATR-IR spectra of unstrained fibrin

**Table S2:** Fitting parameters for ATR-IR spectra of 100% unstrained fibrin

**Figure S3:** tPA binding is reduced to on strained single fibrin fibers

**Figure S4**: SDS page analysis of platelet integrin and BSA coated microbeads

**Figure S4**: Isolated platelet bright filed micrograph showing well-dispersed individual platelets without any aggregate morphology (Scale bar = 50 µm)

**Figure S5**: Platelet attachment and morphology upon exposure to differently strained fibrin

**Movie S1 and S2**: Mechanical deformation of fibrin hydrogel using home build motorized tensile stretcher and THORLABS motorized actuator

**Movie S3**: Z-stack confocal projections of unstrained fibrin showing distribution of ThT and tPA

**Movie S4**: Z-stack confocal projections of 100% strained fibrin showing distribution of ThT and tPA

**Movie S5**: Z-stack confocal projections of unbinding of tPA upon mechanical straining of fibrin

***Fibrin stretching on thin PDMS sheets***

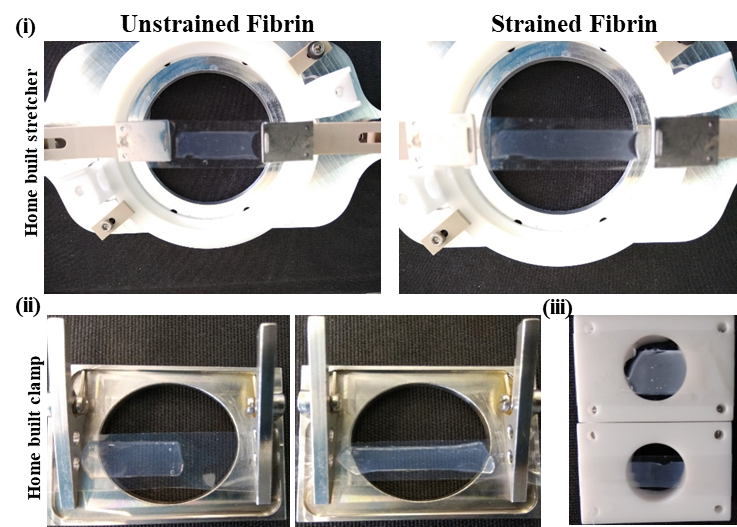

**Figure S1**: Fibrin stretching on thin PDMS sheets. (i) Digital images of initial unstrained and 100% strained fibrin using home-built mechanical stretcher, (ii) and (iii) Clamping of unstrained and stained fibrin gel using home-built clamps.

***Characterization of fibrin structure by ATR-IR spectroscopy***

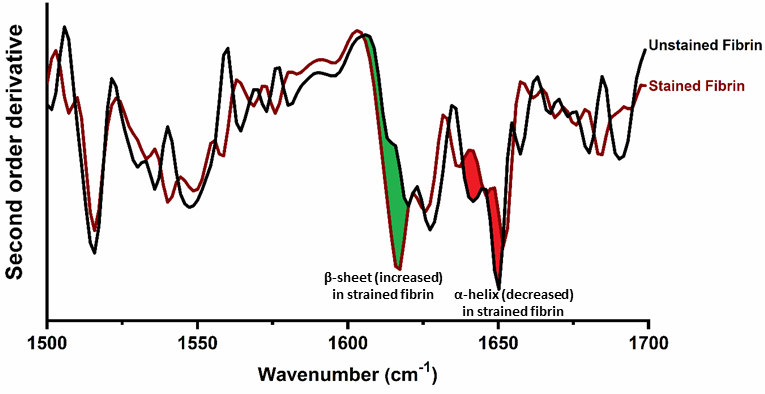

**Figure S2**: Second derivative of IR spectra of unstrained and 100% strained fibrin showing increase in β-sheet content (area highlighted in green) and decrease in α helix content (area highlighted in red) upon mechanical deformation.

**Table S1:** Unstrained Fibrin ATR-IR spectral information

| Peak center (cm^-1^) | FWHM (cm^-1^) | Peak Area (%) | Peak assignment[1-4] |
| --- | --- | --- | --- |
| 1517 | 13 | 4.2 | Random coil |
| 1532 | 20 | 8.9 | β-sheet |
| 1550 | 32 | 19.7 | α-helix |
| 1616 | 21 | 9.6 | β-sheet |
| 1630 | 21 | 17.7 | β-sheet |
| 1644 | 21 | 19.1 | Random coil |
| 1655 | 21 | 14.4 | α-helix |
| 1673 | 20 | 6.4 | turns/loops |

**Table S2:** Strained Fibrin ATR-IR spectral information

| Peak center (cm^-1^) | FWHM (cm^-1^) | Area (%) | Peak assignment[1-4] |
| --- | --- | --- | --- |
| 1519 | 13 | 4.4 | Random coil |
| 1536 | 21 | 10.2 | β-sheet |
| 1554 | 32 | 17.7 | α-helix |
| 1618 | 29 | 26.0 | β-sheet |
| 1632 | 21 | 13.6 | β-sheet |
| 1645 | 18 | 12.1 | Random coil |
| 1655 | 16 | 8.6 | α-helix |
| 1667 | 14 | 5.0 | turns/loops |
| 1679 | 11 | 2.4 | turns/loops |

***tPA binding to strained single fibrin fibers***

In order to evaluate binding and unbinding of tPA to fibrin, we performed experiments on the single fiber level for unstrained and strained fibers. Single fibrin fibers were seeded on a home‑built caliper stretcher. A low concentration fibrin preparation (0.25 mg/ml; having fibrinogen:fibrinogen-488 (90:10)) was prepared on supporting PDMS blocks as described previously and shown in **Figure S3(i-iii)**. A sparse fibrin network was supported on PDMS blocks (**Figure S3(ii)**) and incubated with tPA (Texas red-conjugated tPA, Abcam) protein solution as described in the Methods section of the main text. Confocal microscopy shows that tPA bound to unstrained fibrin between the two PDMS support blocks and revealed a network architecture similar to that seen with fluorescent fibrin (**Fig. S3(iv, top)**). Using the caliper, the PDMS blocks were stretched apart by 100% of their initial separation which stretched the fibrin gel by the same amount (**Fig. S3(iii)**). This was done on the microscope, so the sample could be imaged directly with confocal microscopy. All confocal images were captured under identical operating conditions.

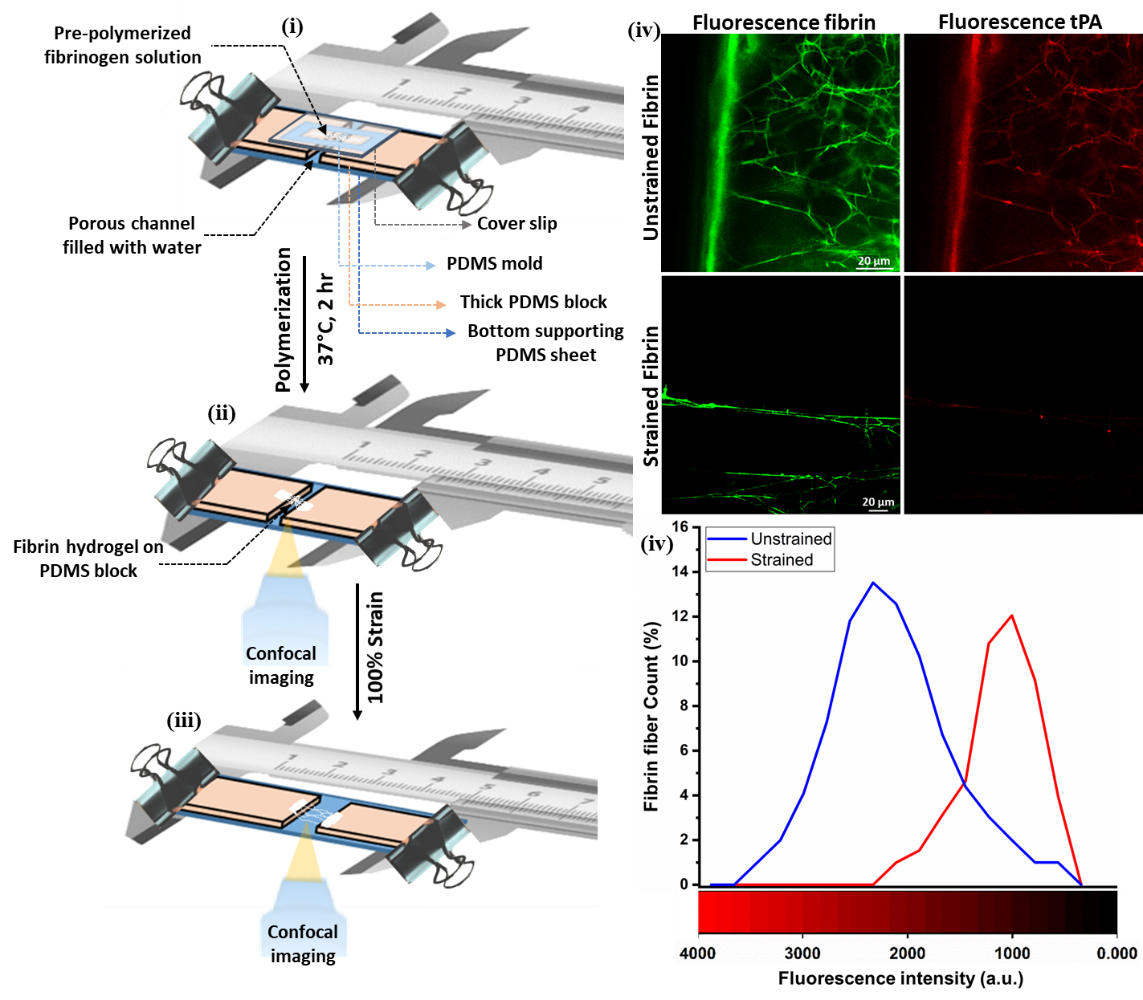

**Figure S3:** tPA binding is reduced to on strained single fibrin fibers. (i-iii) Schematic illustration of preparation of fibrin hydrogel between two PDMS blocks using a caliper stretcher for confocal imaging. (iv) Confocal micrographs of tPA bound to unstrained and strained individual fibers fibrin and (v) tPA fluorescence intensity distribution for unstrained and strained individual fibrin fibers (n ~ 30).

Upon tensile deformation of individual fibers, the intensity of bound fluorescence tPA significantly decreased (**Fig. S3(iv, bottom)**). **Figure S3(v)** shows tPA fluorescence intensity distribution for unstrained and strained individual fibers (n ~ 30). Mean tPA fluorescence shifted from 2200 on unstrained to 1000 on 100% strained fibrin, showing that tPA binding is reduced on stretched fibrin fibers.

***Characterization of bound integrin proteins to the microbeads***

Sodium dodecyl sulfate (SDS) polyacrylamide gel electrophoresis (PAGE) was employed to characterize bound protein (integrin and BSA) on microbeads. Proteins coated microbeads were loaded into sample buffer (NuPAGE LDS) on 4–12% gradient gels and run against standard integrin and BSA as control along with high-molecular-weight marker (ranging from 10 to 225 kDa (Novagen)) as described previously [5].

**Figure S4** shows a Coomassie stained SDS PAGE gel of integrin and BSA proteins from coated microbeads. Integrin and BSA protein bands from beads were compared with unattached integrin and BSA as controls. Lane 1 with pristine beads alone showed no protein, as expected. . Lane 2 for commercial BSA-FITC showed presence of strong protein bands around 62 kDa, Lane 3 showed BSA from BSA-coated beads, which was similar to the BSA control band (Lane 2). Control integrin (α_IIb_β_3_) in lane 4 showed presence of two bands round 100 kDa. As integrin (α_IIb_β_3_) is composed of two different protein chain (α_IIb_) and (β_3_) which are linked by disulfide bond. Under reducing conditions for SDS PAGE, integrin (α_IIb_β_3_) separates into two subunit bands at 125 kDa and 108 kDa for α_IIb_ and β_3_, respectively [6, 7]. Lane 5 shows beads coated with BSA-FITC, which is identical to Lanes 2 and 3. Similarly, in Lane 6, beads coated with BSA and integrin showed presence of specific protein bands for BSA at 62 kDa and for integrin two subunits as shown by arrows.

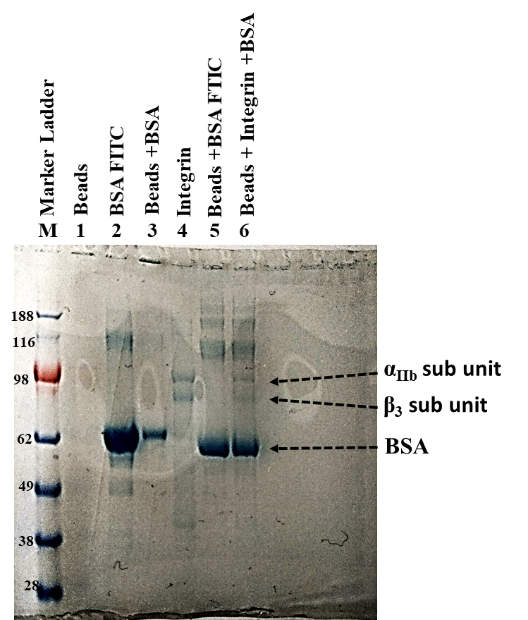

**Figure S4**: SDS page analysis of platelet integrin and BSA coated microbeads. Digital image of SDS PAGE gel for protein-coated beads used for platelet integrin experiments.

***Platelet preparation and characterization on fibrin networks***

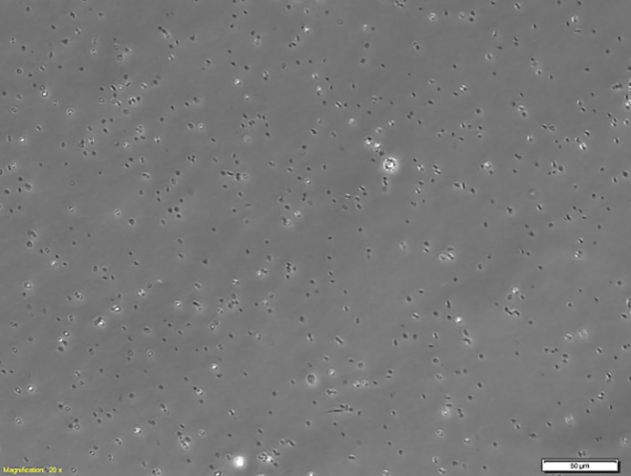

**Figure S5**: Isolated platelet morphology from brightfield images showing well-dispersed individual platelets without any aggregate morphology (scale bar = 50 µm).

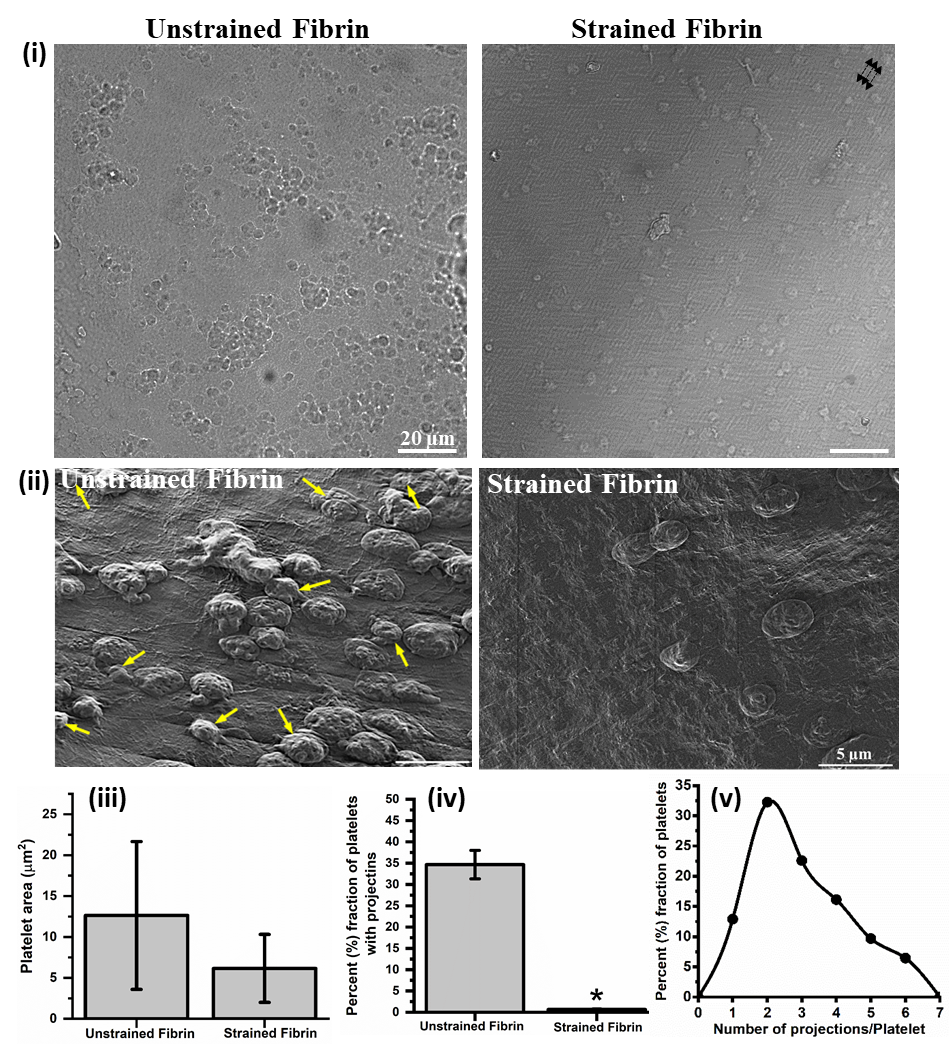

**Figure S6**: Platelet attachment and morphology upon exposure to differently strained fibrin. (i) Bright field images of attached platelets on unstrained and 100% strained fibrin (black arrows indicate direction of fibrin stretch). (ii) SEM micrograph of platelets on unstrained and strained fibrin surface (arrow indicate platelets with projections). (iii) Quantification of platelet spreading (surface area) on unstrained and strained fibrin hydrogel. (iv) and (v) Fraction of attached platelets with multiple projections and distribution profile of number of projections per platelets on unstrained fibrin * indicated statistically significant differences w.r.t. unstrained fibrin based on a t-test with P <0.05.
